# Acute characterization of tissue and functional deficits in a clinically translatable pig model of ischemic stroke

**DOI:** 10.1101/740159

**Authors:** Erin E. Kaiser, Elizabeth S. Waters, Madison M. Fagan, Kelly M. Scheulin, Simon R. Platt, Julie H. Jeon, Xi Fang, Holly A. Kinder, Soo K. Shin, Kylee J. Duberstein, Hea J. Park, Franklin D. West

**Affiliations:** Regenerative Bioscience Center, University of Georgia, Athens, Georgia, United States of America; Neuroscience Program, Biomedical and Health Sciences Institute, University of Georgia, Athens, Georgia, United States of America; Department of Animal and Dairy Science, College of Agricultural and Environmental Sciences at the University of Georgia, Athens, Georgia, United States of America; Department of Small Animal Medicine and Surgery, College of Veterinary Medicine at the University of Georgia, Athens, Georgia, United States of America; Department of Foods and Nutrition, College of Family and Consumer Sciences at the University of Georgia, Athens, Georgia, United States of America; Department of Pharmaceutical and Biomedical Sciences, Interdisciplinary Toxicology Institute, University of Georgia, Athens, Georgia, United States of America

**Keywords:** brain ischemia, gait analysis, magnetic resonance imaging, porcine, acute stroke

## Abstract

The acute stroke phase is a critical time frame used to evaluate stroke severity, therapeutic options, and prognosis while also serving as a major target for the development of diagnostics. To better understand stroke pathophysiology and to enhance the development of treatments, our group developed a translational pig ischemic stroke model. In this study, the evolution of acute ischemic stroke tissue damage, immune response, and functional deficits were further characterized in the pig model. Stroke was induced by middle cerebral artery occlusion in Landrace pigs. At 24 hours post-stroke, magnetic resonance imaging revealed a decrease in ipsilateral diffusivity and an increase in hemispheric swelling and intracranial hemorrhage resulting in notable midline shift. Stroke negatively impacted white matter integrity leading to decreased fractional anisotropy. Similar to acute clinical patients, stroked pigs showed a reduction in circulating lymphocytes and a surge in neutrophils and band cells. Functional responses corresponded with structural changes with reduced exploration in open field testing and impairments in spatiotemporal gait parameters. This novel, acute ischemia characterization provides important insights into tissue and functional level changes in a pig model that can be used to identify treatment targets and future testing of therapeutics and diagnostics.

## Introduction

Every year, 6.2 million people worldwide die from stroke making it the leading cause of death in individuals over the age of 60 and the fifth leading cause of death in individuals ages 15-59 (1, 2). Of the patients that survive, approximately 5 million are left permanently disabled making stroke a global medical and socioeconomic problem (3). The acute phase of ischemic stroke is a critical time window to determine stroke severity, treatment options, and future prognosis in clinical patients. Specifically, the acute phase is a major target for the development of novel therapeutics and diagnostics as an early reduction in brain tissue loss is directly correlated with improvements in functional outcomes. In addition, all current Food and Drug Administrative approved treatments, tissue plasminogen activator (tPA) and thrombectomy, are only effective during this acute window (4–6). The acute phase of ischemic stroke has also been the focus of diagnostic and prognostic tool development; tools including magnetic resonance imaging (MRI) that can rapidly and accurately identify ischemic stroke and has demonstrated strong predictive value with respect to long-term patient outcomes (7–11). However, the development of therapies and diagnostic tools has been slower than desired particularly with respect to treatments with numerous failed clinical trials (12–15).

A potential opportunity to hasten the speed at which therapies and diagnostics reach patients is through the use of translational large animal models that are more predictive of human outcomes. Assessments by the Stem Cell Emerging Paradigm in Stroke (STEPS) and the Stroke Therapy Academic Industry Roundtable (STAIR) consortiums identified therapeutic testing in higher-order gyrencephalic species and in translational animal models more reflective of human pathology and physiology as major needs in pre-clinical stroke studies to better predict therapeutic efficacy (6, 16–21). To address this unmet need, a pig ischemic stroke model has been recently developed by our research team with anatomy, physiology, and stroke pathology similar to human patients (22–25). The pig brain is similar in mass compared to humans being only 7.5 times smaller, whereas the rodent brain is 650 times smaller in comparison to humans (26). This allots for a more direct assessment of therapeutic dosing in a pre-clinical model. The pig’s brain size is also an advantage in developing diagnostic tools as human 3T MRI scanners and coils can be used to develop new MRI sequences and analytical tools. In terms of cytoarchitecture, human and pig brains are gyrencephalic and are composed of >60% white matter (WM), while rodent brains are lissencephalic and are composed of <10% WM, making pig tissue responses potentially more predictive of human outcomes (27–30). These attributes are critically important as WM and gray matter (GM) exhibit differing sensitivities to hypoxia (30). Although the failure of pharmacological translation is multifactorial, the failure to ameliorate ischemic damage to WM is proposed to be a major factor (31). The similarities between pig and humans in brain size, cytoarchitecture, and WM composition collectively support the use of a pig ischemic stroke model to more accurately predict potential outcomes of human clinical trials. However, more in depth characterization of the acute ischemic stroke timepoint is needed in the pig model to better understand similarities and differences between human and pig acute stroke outcomes.

MRI is an excellent tool for use in the pig ischemic stroke model as it allows for bidirectional development of the pig model as well as MRI diagnostic and prognostics. MRI allows for the assessment of stroke evolution in the pig model and evaluation of novel therapeutic efficacy. In addition, new MRI sequences and post-processing tools can be developed in the pig for use in clinical settings. Acute MRI assessment of ischemic stroke patients has become the standard of care in diagnosing and predicting patient clinical outcomes (8, 32). Clinically, diffusion-weighted imaging (DWI) has been shown to reliably enable early identification of the lesion size, location, and age with high sensitivity and specificity (7, 33–38). Moreover, acute stage DWI lesion volume measures have proven to be highly correlated with chronic lesion size and stroke severity as determined by Modified Rankin Scale (mRS) and National Institutes of Health Stroke Scale (NIHSS), suggesting DWI provides valuable prognostic information (7-9, 38-40). DWI derived apparent diffusion coefficient (ADC) maps have aided in further understanding the time course of acute ischemic brain damage by tracking the diffusion of water in the hypoxic brain parenchyma from extracellular to intracellular compartments (41, 42). In conjunction with other MR techniques, ADC hypointensities allow clinicians to differentiate between regions at risk for cerebral infarction and irreversibly damaged tissue in order to establish time windows for stroke treatment and to identify patients who are most likely to benefit from acute stroke therapies (7, 40, 43, 44). Disruption of WM structural integrity is also associated with poor early neurological outcomes in stroke patients (45). Diffusion tensor imaging (DTI) studies of human stroke reveal notable alterations in WM fractional anisotropy (FA) that correspond with the temporal evolution of stroke (10, 11). FA analysis has improved the identification of ischemic lesions at acute and subacute time points and has proved particularly useful in determining time of stroke onset, which is frequently unknown in clinical settings (11). Recently, progressive structural remodeling of contralateral WM tracts related to motor, cognitive, and sensory processing was positively associated with motor function recovery in the acute and sub-acute stages post-stroke as well as 1, 4, and 12 weeks post-ischemic onset in patients (46, 47). Acute MRI analysis in the pig stroke model will allow for the characterization of clinically relevant parameters and to assess for correlations with acute functional changes as observed in human patients.

Ischemic stroke leads to a wide array of acute deficits in behavior, cognition, and sensorimotor function in clinical patients thus resulting in poor mRS scale scores (48). Neurological deficits in executive function, episodic memory, visuospatial function, and language manifest within 48 to 72 hours in 33.6% of patients (49–52). Occlusions of the middle cerebral artery (MCA) and territorial infarction are regularly linked to acute limb paresis that is sustained long-term (52). Understanding these motor impairments are essential to planning rehabilitation efforts to restore ambulatory activity levels and balance deficiencies in stroke survivors (53, 54). Specifically, improvements in foot placement, stride length, and walking speed are recognized as powerful indicators of long-term recovery (55–59). Among these neurologic and functional consequences, post-stroke depression (PSD) is the most frequent psychiatric problem occurring in one-third of stroke survivors (60). PSD is strongly associated with further inhibition of recovery processes due to the combination of ischemia-induced neurobiological dysfunctions and psychosocial distress (61, 62). The pig stroke model offers a unique opportunity to study acute changes in behavior, cognition, and motor function due to anatomical similarities in the size of the prefrontal cortex and cerebellum in addition to somatotopical organization of the motor and somatosensory cortices which are critically important in modeling human motor function effects in the acute ischemic stroke phase (26, 63–65).

The objective of this study was to utilize clincially relevant assessment modalities to characterize acute ischemic stroke in a pig model that will provide a translational platform to study potential diagnostics and therapeutic interventions. We present evidence pigs display an acute ischemic stroke response similar to human patients including brain lesioning, swelling, loss of WM integrity, and increased white blood cell (WBC) counts. These physiological changes correlated with aberrant behavior and worsened motor function. This compelling evidence suggests the pig stroke model could serve as a valuable tool in bridging the gap between pre-clinical rodent studies and human clinical trials.

## Materials and methods

### Animals and housing

All work performed in this study was approved by the University of Georgia Institutional Animal Care and Use Committee (IACUC; Protocol Number: 2017-07-019Y1A0) and in accordance with the National Institutes of Health Guide for the Care and Use of Laboratory Animals guidelines. 6, sexually mature, castrated male Landrace pigs, 5-6 months old and 48-56 kg were purchased from the University of Georgia Swine Unit and enrolled in this study. Male pigs were used in accordance with the STAIR guidelines that suggests initial therapeutic evaluations should be performed with young, healthy male animals (66). Pigs were individually housed in a Public Health Service (PHS) and AAALAC approved facility at a room temperature approximately 27°C with a 12 hour light/dark cycle. Pigs were given access to water and fed standard grower diets with provision of enrichment through daily human contact and toys.

### Study design

The sample size for this study was determined by a power calculation based on our routine use of the middle cerebral artery occlusion model with lesion volume changes by MRI imaging being the primary endpoint (67). The power analysis was calculated using a two-tailed ANOVA test, α=0.05, and an 80% power of detection effect size of 1.19 and a standard deviation of 44.63. This was a randomized study where 2 pigs were randomly assigned to each surgical day. All endpoints and functional measurements were prospectively planned and underwent blinded analysis. Predefined exclusion criteria from all endpoints included instances of infection at the incision site, self-inflicted injuries that required euthanasia, inability to thermoregulate, uncontrolled seizure activity, and/or respiratory distress. 1 pig was excluded from MRI collection as well as post-stroke blood and functional analysis due to post-operative complications and premature death. No outliers were removed from the data.

### Middle cerebral artery occlusion surgical procedures

The day prior to surgery, pigs were administered antibiotics (Excede; 5 mg/kg intramuscular (IM) and fentanyl for pain management (fentanyl patch; 100 mg/kg/hr transdermal (TD)). Pre-induction analgesia and sedation were achieved using xylazine (2 mg/kg IM) and midazolam (0.2 mg/kg IM). Anesthesia was induced with intravenous (IV) propofol to effect and prophylactic lidocaine (1.0 mL 2% lidocaine) topically to the laryngeal folds to facilitate intubation. Anesthesia was maintained with isoflurane (Abbott Laboratories) in oxygen.

As previously described, a curvilinear skin incision extended from the right orbit to an area rostral to the auricle (24). A segment of zygomatic arch was resected while the temporal fascia and muscle were elevated and a craniectomy was performed exposing the local dura mater. The distal middle cerebral artery (MCA) and associated branches were permanently occluded using bipolar cautery forceps resulting in ischemic infarction. The temporalis muscle and epidermis were routinely re-apposed.

Anesthesia was discontinued and pigs were returned to their pens upon extubation and monitored every 15 minutes until vitals including temperature, heart rate, and respiratory rate returned to normal, every 4 hours for 24 hours, and twice a day thereafter until post-transplantation sutures were removed. Banamine (2.2 mg/kg IM) was administered for post-operative pain, acute inflammation, and fever management every 12 hours for the first 24 hours, and every 24 hours for 3 days post-stroke.

### Magnetic resonance imaging acquisition and analysis

MRI was performed 24 hours post-stroke on a General Electric 3.0 Tesla MRI system. Pigs were sedated and maintained under anesthesia as previously described for MCAO surgery. MRI of the cranium was performed using an 8-channel torso coil with the pig positioned in supine recumbency. Multiplanar MR brain imaging sequences were acquired including T2 Fluid Attenuated Inversion Recovery (T2FLAIR), T2Weighted (T2W), T2Star (T2*), DWI, and DTI. Sequences were analyzed using Osirix software. Cytotoxic edema consistent with ischemic stroke was confirmed 24 hours post-stroke by comparing corresponding hyperintense regions in T2FLAIR and DWI sequences and hypointense regions in ADC maps.

DWI sequences were used to generate ADC maps. ADC values were calculated for each axial slice at a manually drawn region of interest (ROI) that was defined by areas of hypointensity and directly compared to an identical ROI in the contralateral hemisphere. Average ADC values were obtained by calculating the average signal intensity across all slices and reported as 10^-3^ mm^2^/s. Hemisphere volume was calculated using T2W sequences for each axial slice by manually outlining the ipsilateral and contralateral hemispheres. The hemisphere areas were multiplied by the slice thickness (3mm) to obtain total hemisphere volumes. Lesion volume was calculated using DWI sequences for each axial slice by manually outlining hyperintense ROIs. The area of each ROI was multiplied by the slice thickness (2mm) to obtain the total lesion volume. Similarly, intracranial hemorrhage (ICH) volume was calculated by manually outlining areas of hypointensity utilizing T2* sequences. Midline shift (MLS) was calculated utilizing T2W sequences for each axial slice by measuring the distance from the natural midline along the anterior and posterior attachments of the falx cerebri to the septum pellucidum. DTI was utilized to generate FA maps. FA values of the internal capsules were calculated manually on one representative slice per pig and were expressed as a percent change in the ipsilateral hemisphere relative to the contralateral hemisphere.

### Blood collection and analysis

Venous blood samples were collected pre-stroke, 4, 12, and 24 hours post-stroke into K2EDTA spray coated tubes (Patterson Veterinary). 4 μL of blood was pipetted onto the base of a ColorFrost microscope slide (ThermoScientific) approximately 1 cm from the edge. At an angle of approximately 45 degrees, a spreader slide was placed in front of the blood and retracted until the blood sample evenly spread along the width of the slide. Even pressure on the spreader slide was applied in a forward direction in order to create a smear. Care was taken to ensure each blood smear covered two-thirds of the slide and exhibited an oval feathered end. Each slide was air dried for 10 minutes, fixed with methanol for 2 minutes, air dried for 2 minutes, and then stained in Wright-Giemsa stain for 5 minutes. The stained slide was submerged in distilled water (dH_2_O) for 10 minutes. Finally, the slide was rinsed, air dried, and then a cover slip was applied using Phosphate Buffered Saline (PBS). Trained, blinded personnel completed manual white blood cell counts of lymphocytes, neutrophils, and band cells at the monolayer, beginning approximately one millimeter away from the body of the smear. The first 100 white blood cells visualized were identified and cell counts were expressed as a percentage.

### Gait analysis

Pigs underwent gait analysis pre-stroke and 48 hours post-stroke to assess changes in spatiotemporal gait parameters. Data was recorded using a GAITFour^®^ electronic, pressure-sensitive mat (CIR Systems Inc., Franklin, NJ) 7.01 m in length and 0.85 m in width with an active area that is 6.10 m in length and 0.61 m in width. In this arrangement, the active area is a grid, 48 sensors wide by 480 sensors long, totaling 23,040 sensors. 2 weeks pre-stroke, pigs were trained to travel across a gait mat at a consistent, 2-beat pace. To reinforce consistency, rewards were given at each end of the mat for successful runs. Pre-stroke gait data was collected on 3 separate days for each pig. At each time point, pigs were encouraged to move along the mat until 5 consistent trials were collected in which the pigs were not distracted and maintained a consistent pace with no more than 15 total trials collected.

Gait data was semi-automatically analyzed using GAITFour^®^ Software to provide quantitative measurements of velocity (cm/sec) and cadence (steps/min). Additional measurements were quantified specifically for the affected front left limb, which is contralateral to the induced stroke lesion on the right side of the brain. These measurements included stride length (the distance between successive ground contact of the same hoof), swing percent of cycle (the percent of a full gait cycle in which a limb is not in contact with the ground), cycle time (the amount of time for a full stride cycle), swing time (the amount of time a limb is in the swing phase, or not in contact with the ground) and mean pressure (the amount of pressure exerted by a limb).

### Open field testing

As an additional measure of functional outcome, pigs underwent open field (OF) behavior testing pre-stroke and 48 hours post-stroke. All tests took place in a 2.7 m x 2.7 m arena lined with black rubber matting, used to provide stable footing. White curtains were hung around the arena to reduce visual distractions during testing. Trials were recorded using EthoVision video tracking software (Noldus Systems) to obtain objective and quantifiable measures of behavioral characteristics.

Pigs were individually brought to the behavior arena and allowed to explore for 10 minutes during the OF test. Behaviors automatically tracked during this test include velocity and distance traveled. Additionally, exploratory behaviors typical of pigs such as sniffing the wall (perimeter sniffing) were manually tracked and coded in the EthoVision software by trained personnel.

### Statistical analysis

All quantitative data was analyzed with SAS version 9.3 (Cary, NC) and statistical significances between groups were determined by one-way analysis of variance (ANOVA) and post-hoc Tukey-Kramer Pair-Wise comparisons. Comparisons where p-values were ≤ 0.05 were considered significantly different.

## Results

### MCAO induces acute ischemic infarction and decreased diffusivity

To confirm ischemic stroke 24 hours post-MCAO, MRI DWI (Fig 1A) and T2FLAIR sequences were assessed. Scans exhibited territorial hyperintense lesions characteristic of an edematous injury. Hypointense lesions observed on corresponding ADC maps (Fig 1B) confirmed areas of restricted diffusion indicative of cytotoxic edema thus confirming permanent cauterization of the MCA resulted in ischemic stroke. DWI-ADC mismatch resulted in identification of potentially salvageable penumbra tissue. DWI sequences revealed an average lesion volume of 9.91±1.40 cm^3^ (Fig 1A). ADC sequences revealed significantly (p≤0.0001) decreased diffusivity within ischemic lesions when compared to identical regions of interest in the contralateral hemisphere (0.34±0.02 vs. 0.62±0.03 x10^-3^mm^2^/s, respectively; Fig 1B-C).

**Figure 1:**
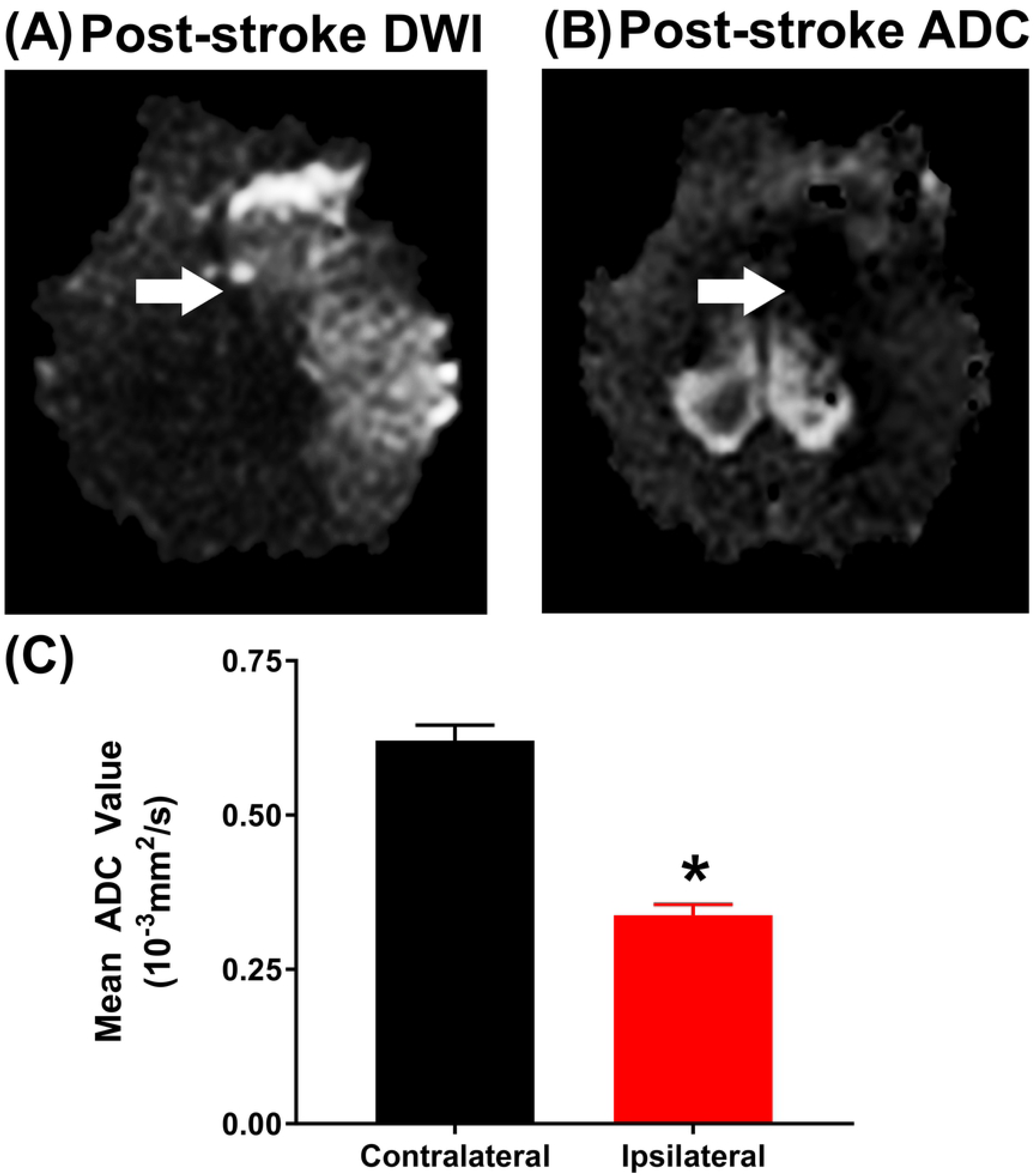
MCAO induces acute ischemic infarction and decreased diffusivity. DWI sequences exhibited territorial hyperintense lesions of 9.91±1.40 cm^3^ characteristic of an edematous injury (A, white arrow). ADC maps revealed signal void indicative of restricted diffusion and cytotoxic edema (B, white arrow). Ipsilateral ROIs exhibited a significantly (p≤0.0001) lower ADC value relative to the contralateral hemisphere (0.34±0.02 vs. 0.62±0.03 x10^-3^mm^2^/s, respectively; C). * indicates significant difference between hemispheres.

### Ischemic stroke results in acute hemispheric swelling, hemorrhage, and loss of white matter integrity

Analysis of T2W sequences at 24 hours post-stroke revealed a trending (p=0.16) increase in ipsilateral hemisphere volume indicative of cerebral swelling when compared to the contralateral hemisphere (25.99±1.78 vs. 22.49±1.40 cm^3^, respectively; Fig 2A-C) and an associated MLS of 2.48±0.55 mm (Fig 2A-B). Acute ICH was observed via T2* sequences with a consistent mean hemorrhage volume of 1.73±0.07 cm^3^ (Fig 2D-E, white arrow), which suggests the ischemic infarct area underwent hemorrhagic transformation (HT). These HTs impacted basal ganglion structures as well as portions of the cerebellum, brain regions responsible for motor function. To assess changes in WM integrity, FA values of the internal capsules were evaluated 24 hours post-stroke, revealing a significant (p<0.01) decrease in the ipsilateral internal capsule (IC) when compared to the contralateral side (0.17±0.01 vs. 0.23±0.01 respectively; Fig 3A-C). Collectively, MRI results demonstrated MCAO led to tissue-level damage including ischemic infarction, decreased diffusivity, hemispheric swelling, pronounced MLS, HT, and disrupted WM integrity.

**Figure 2:**
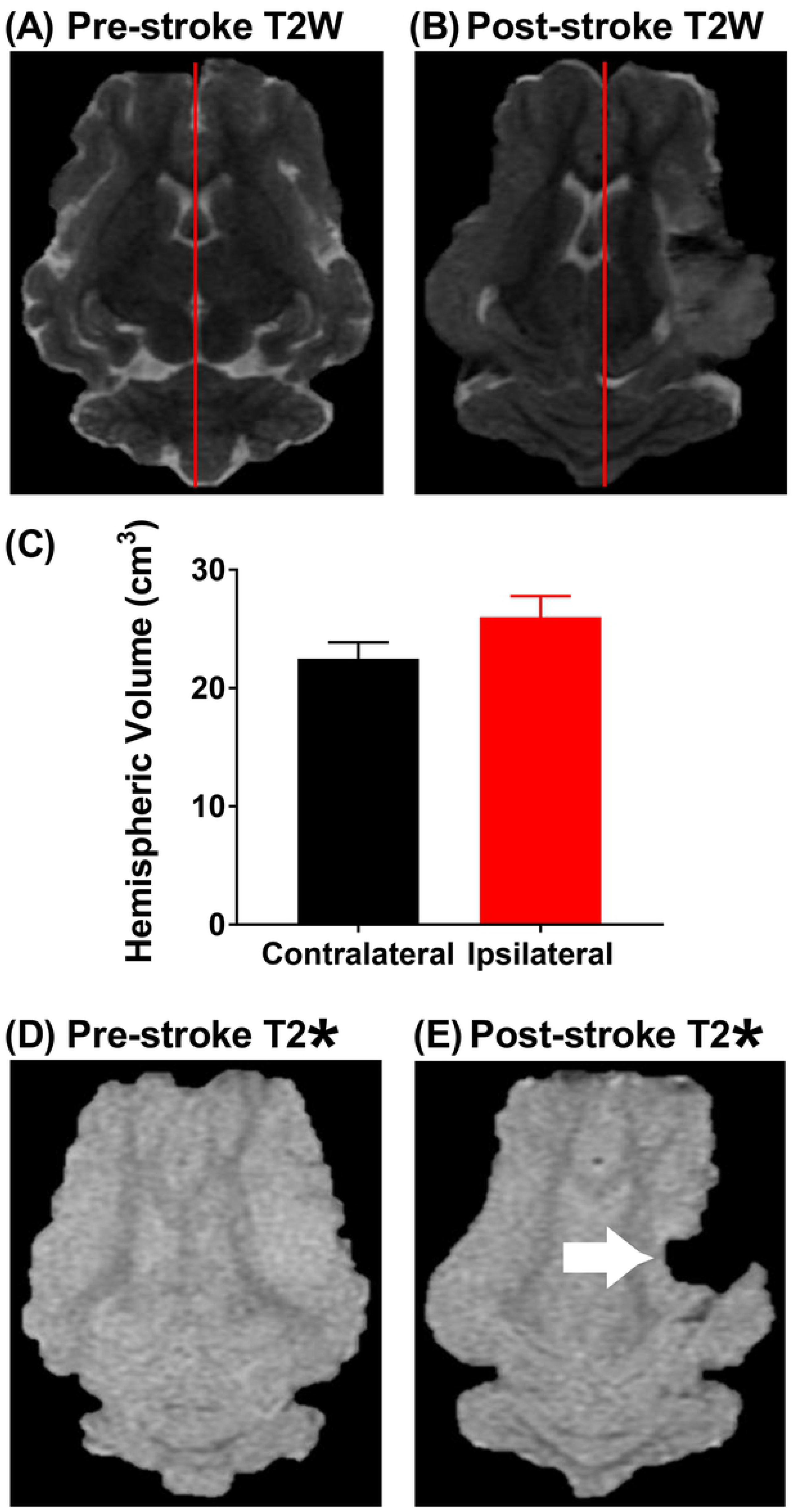
Ischemic stroke results in hemispheric swelling, consequent midline shift, and intracranial hemorrhage. T2W sequences revealed increased swelling of the ipsilateral hemisphere (25.99±1.78 vs. 22.49±1.40 cm^3^; A-C) resulting in a pronounced MLS of 2.48±0.55 mm compared to pre-stroke imaging (A and B, red lines). Characteristic hypointense ROIs indicated the presence of ipsilateral ICH when compared to pre-stroke T2* sequences (1.73±0.17 cm^3^, D and E, white arrow).

**Figure 3:**
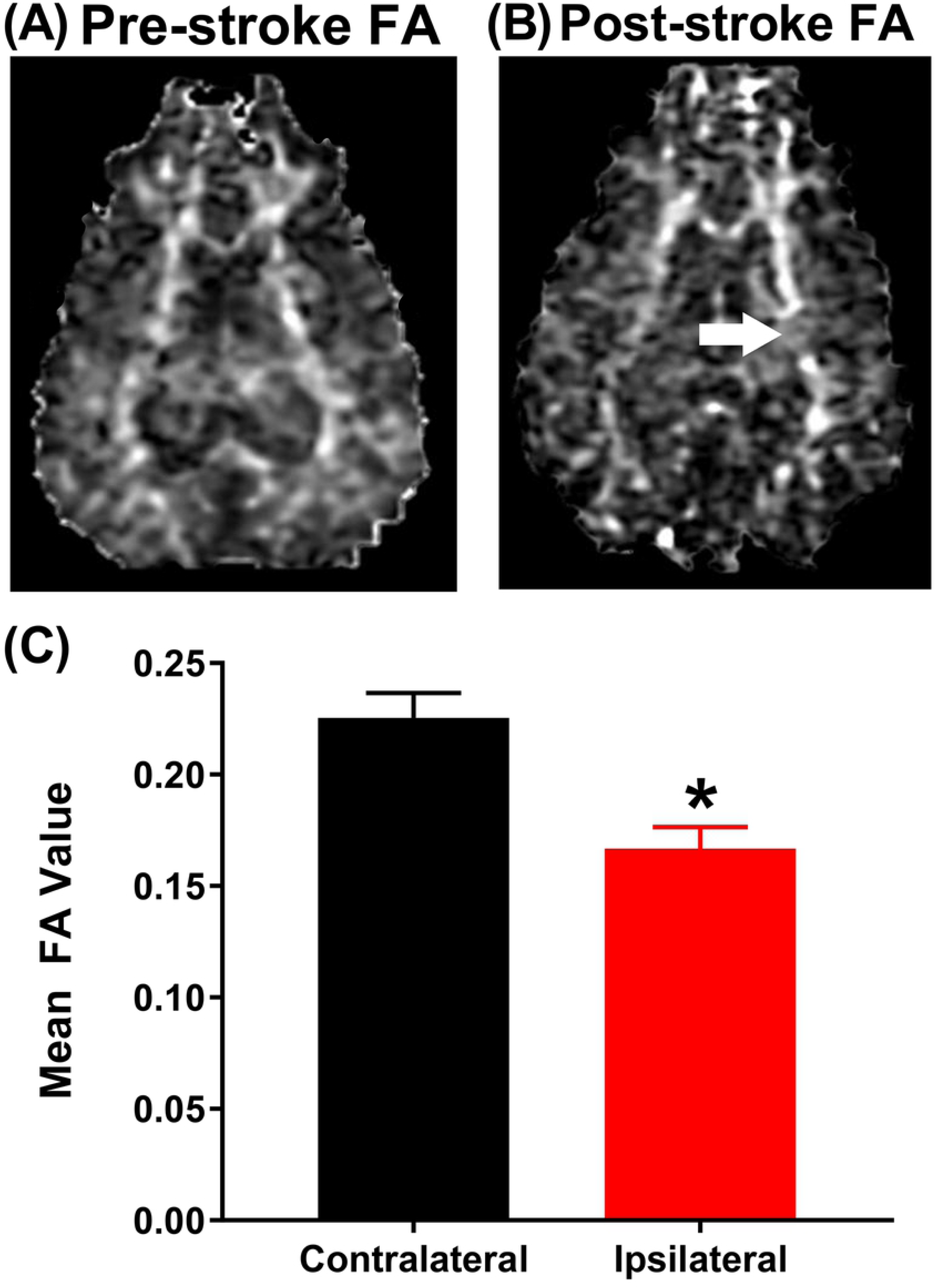
Ischemic stroke diminishes white matter integrity of the internal capsule. Pre-stroke the left and right IC possess similar WM integrity (A). 24 hours post-stroke, the ipsilateral IC exhibited a disruption in WM integrity (B, white arrow). Further analysis revealed a significant (p<0.01) decrease in the ipsilateral IC FA value when compared to the contralateral IC (0.17±0.01 vs. 0.23±0.01 respectively; C). * indicates significant difference between hemispheres.

### Ischemic stroke increases circulating neutrophil levels and decreases circulating lymphocyte levels

To determine changes in immune cell response to acute ischemic stroke, venous blood samples were collected pre-stroke, 4, 12, and 24 hours post-stroke. Band neutrophils (Fig 4A-B), neutrophils (Fig 4C-D), and lymphocytes (Fig 4E-F) were assessed via manual cell counts. Band neutrophils significantly (p<0.05) increased 12 hours post-stroke compared to pre-stroke (5.50±0.99% vs. 1.92±0.51% respectively; Fig 4B). Similarly, the number of circulating neutrophils was significantly (p<0.05) increased at 4 and 12 hours post-stroke when compared to pre-stroke (43.7±5.27% and 48.9±3.92% vs. 26.5±1.96%, respectively; Fig 4D). The number of circulating lymphocytes was significantly (p<0.05) decreased at 12 and 24 hours post-stroke compared to pre-stroke (25.60±4.01% and 26.60±4.29% vs. 44.83±3.66% respectively; Fig 4F). These results demonstrated stroke resulted in an increase in circulating band neutrophils and neutrophils and a decrease in circulating lymphocytes which indicates an acute immune response.

**Figure 4:**
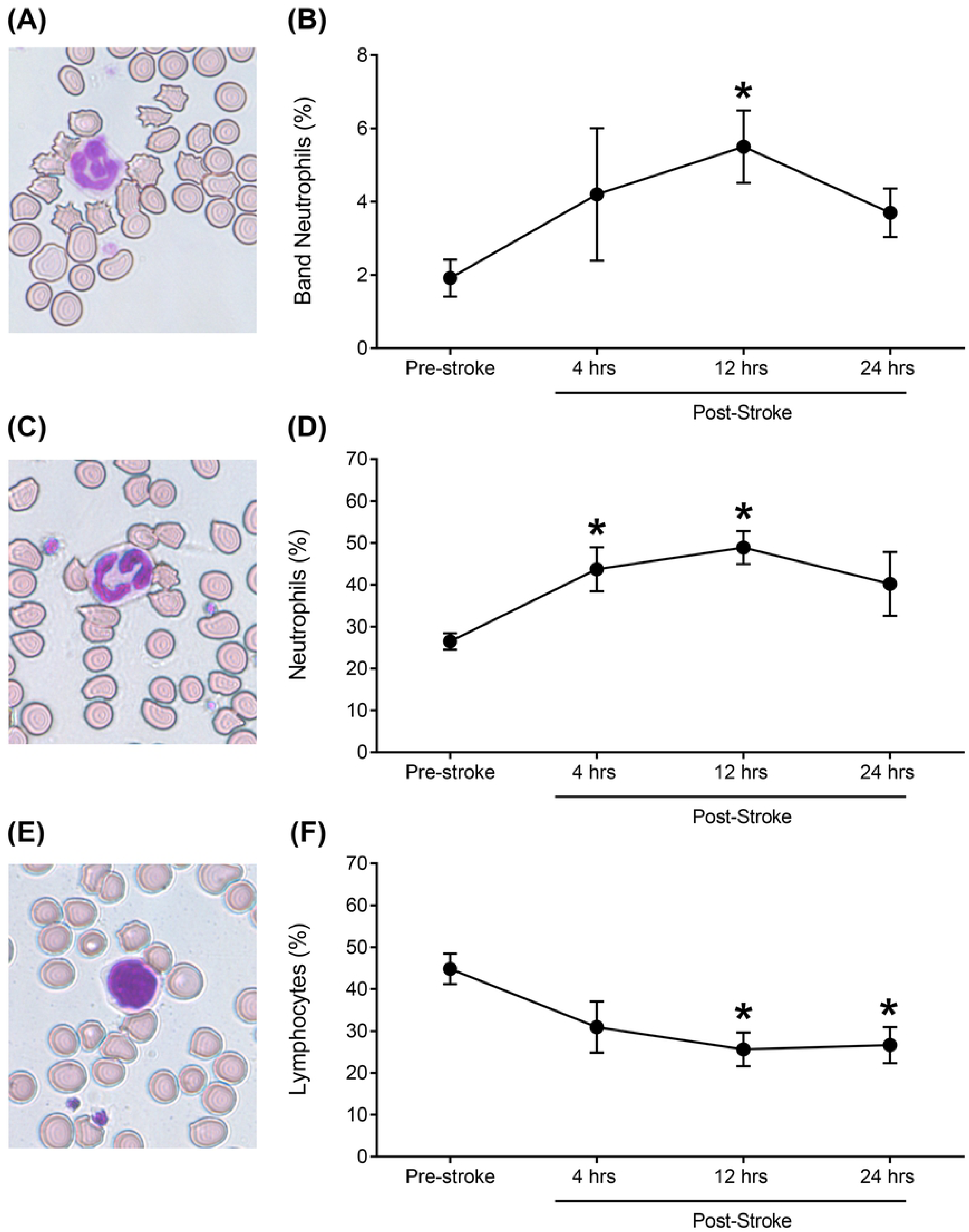
Ischemic stroke leads to increases in circulating neutrophil levels and decreases in circulating lymphocyte levels. Band neutrophils showed a significant (p<0.05) increase 12 hours post-stroke when compared to pre-stroke (5.50±0.99 vs. 1.92±0.51% respectively; A, B). Circulating neutrophils were significantly (p<0.05) increased at 4 and 12 hours post-stroke relative to pre-stroke (43.7±5.27 and 48.9±3.92% vs. 26.5±1.96%, respectively; C, D). Circulating lymphocytes were significantly (p<0.05) decreased at 12 and 24 hours post-stroke compared to pre-stroke (25.60±4.01 and 26.60±4.29% vs. 44.83±3.66% respectively; E, F). * indicates significant difference between pre-stroke and post-stroke time points.

### Ischemic stroke decreases exploratory behaviors during open field testing

Changes in exploratory behaviors were assessed using the open field (OF) test 48 hours post-stroke. Perimeter sniffing, a typical exploratory behavior exhibited by pigs, was recorded utilizing Ethovision XT tracking software to assess differences in perimeter sniffing pre- and post-stroke (Fig 5A-B); representative 10 minute movement tracings show perimeter sniffing (red) and non-perimeter sniffing (yellow). Pigs’ perimeter sniffing frequency significantly (p<0.05) decreased 48 hours post-stroke compared to pre-stroke (13±2.94 vs 26±4.02 times, respectively, Fig 5C). However, no significant differences were noted for velocity and distance traveled in the OF test between pre- and 48 hours post-stroke. These results suggest that stroke impairs normal exploratory behaviors.

**Figure 5:**
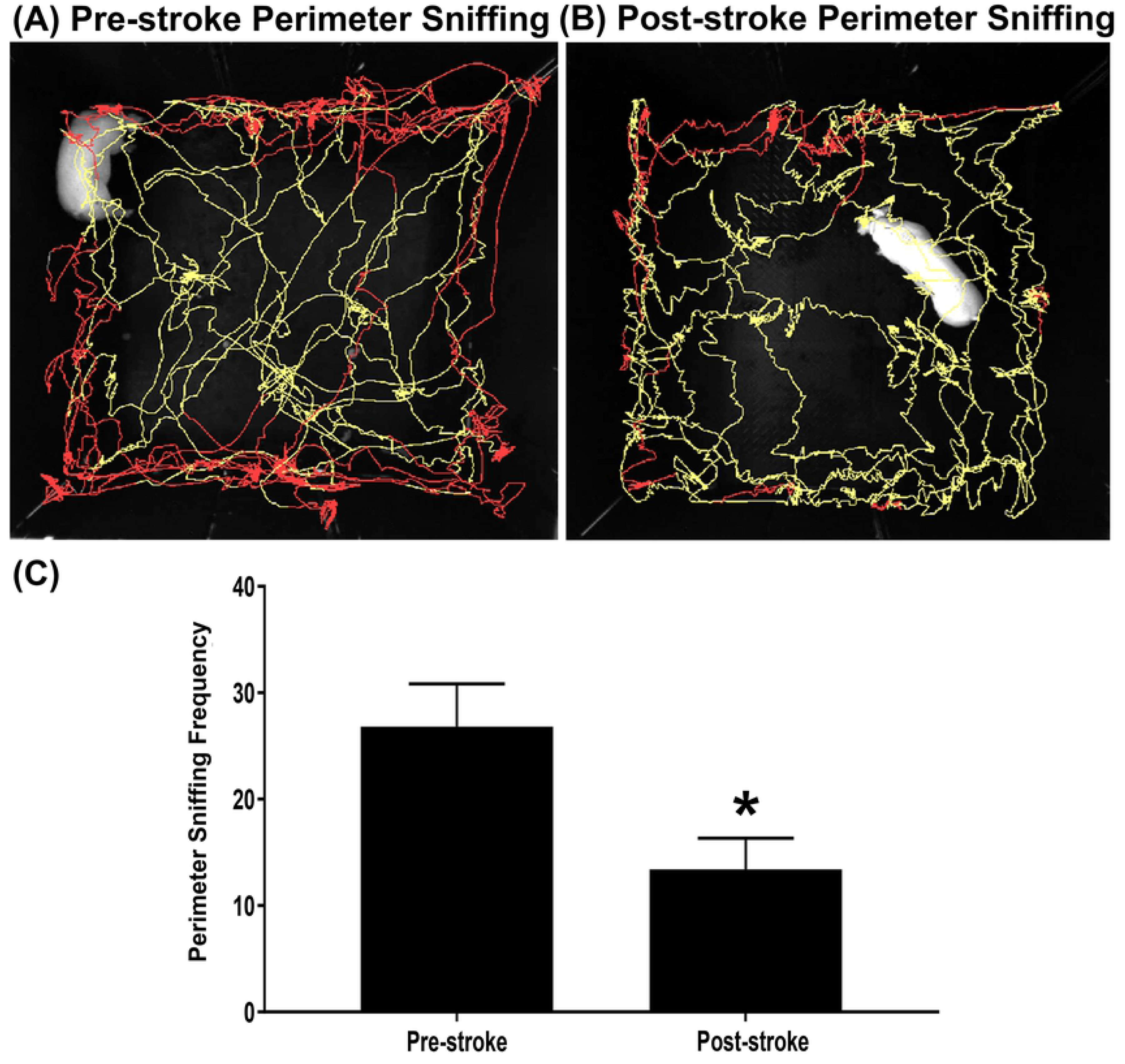
MCAO leads to functional disabilities and behavioral abnormalities. Ethovision XT tracking software was used during OF testing to automatically assess differences in perimeter sniffing (red line) versus OF arena exploration (yellow line) pre-stroke (A) and post-stroke (B). Exploratory perimeter sniffing frequencies were significantly (p<0.05) reduced at 48 hours post-stroke compared to pre-stroke observations (13.0±2.94 vs 26.0±4.02, respectively; C). * indicates a significant difference from pre-stroke.

### Ischemic stroke results in spatiotemporal gait deficits

Key spatiotemporal gait parameters were analyzed pre-stroke and 48 hours post-stroke to detect potential impairments in motor function as an outcome of stroke. Significant (p<0.01) decreases were noted in the average velocity and cadence at 48 hours post-stroke compared to pre-stroke indicating the speed of the pigs decreased as a result of stroke (61.01±8.39 vs 162.9±12.73 cm/s and 61.01±5.91 vs 126.44±3.72 steps/min, respectively, Fig 6A-B). Further changes were noted in measurements of the contralateral left forelimb (LF). The limb contralateral to the stroke lesion typically has more pronounced motor deficits relative to the ipsilateral limb in humans, mice, and rats (68, 69). The swing percent of cycle significantly (p<0.01) decreased demonstrating pigs spent more time with the LF in contact with the ground at 48 hours post-stroke compared to pre-stroke suggesting an increased need for support (30.70±2.12 vs 48.89±2.35%, respectively, Fig 6C). A significant (p<0.01) decrease in stride length of the LF was observed at 48 hours post-stroke compared to pre-stroke (59.04±3.85cm vs 76.72±4.60cm, respectively, Fig 6D). Cycle time of the LF significantly (p<0.01) increased signifying a slower gait at 48 hours post-stroke compared to pre-stroke (1.02±.09 vs 0.48±0.013sec, respectively, Fig 6E). Finally, the mean pressure exhibited by the LF significantly (p<0.01) decreased at 48 hours post-stroke compared to pre-stroke (2.62±.03 vs 2.82±.03 arbitrary units (AU), respectively, Fig 6F). Deficits in the measured gait parameters indicate stroke lead to substantial motor impairments at acute time points in pigs.

**Figure 6:**
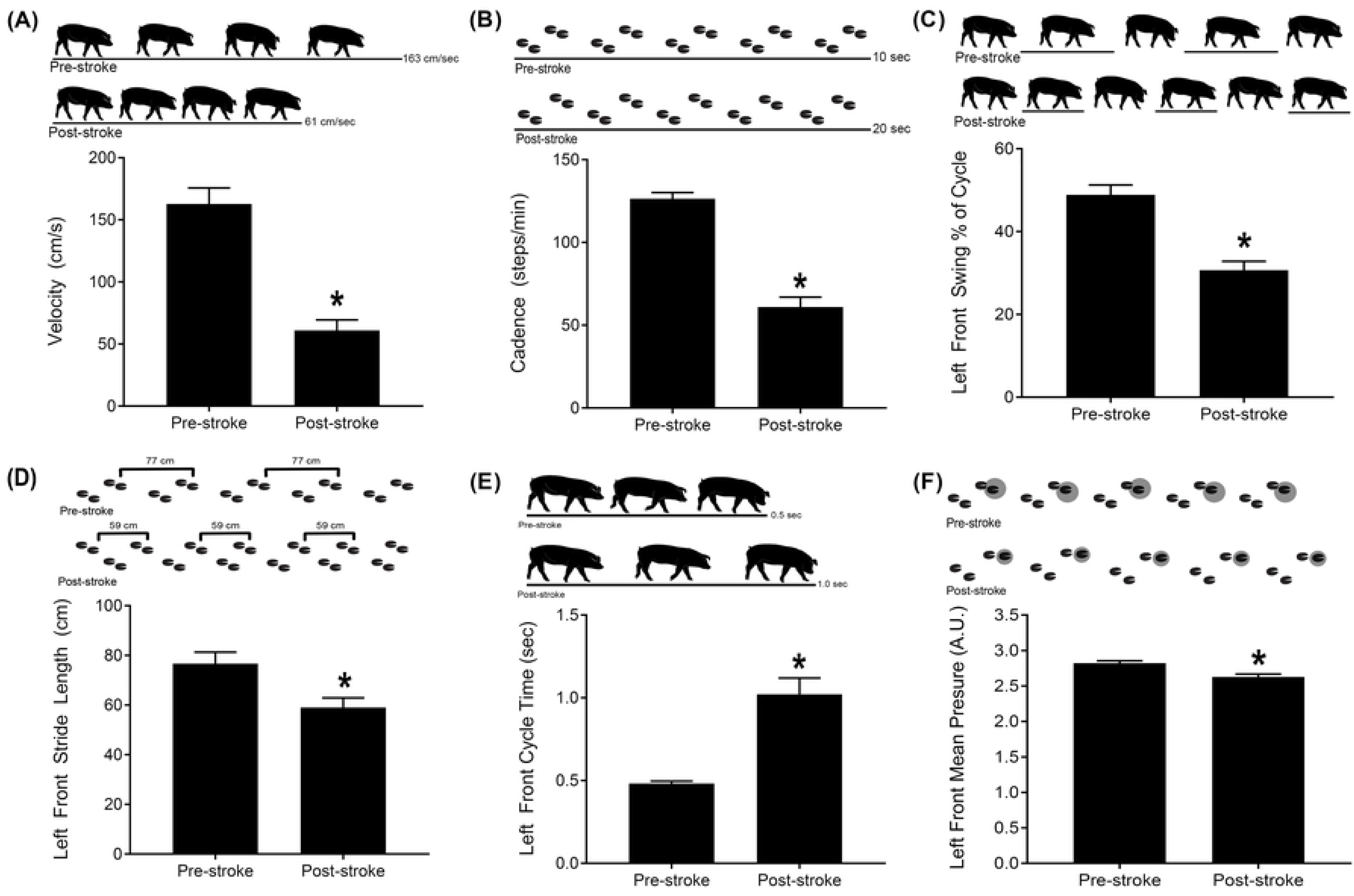
Ischemic stroke results in spatiotemporal gait deficits. Velocity and cadence significantly (p< 0.01) decreased post-stroke (61.01±8.39 vs 162.9±12.73 cm/s and 61.01±5.91 vs 126.44±3.72 steps/min, respectively, A-B). The LF swing percent of cycle significantly (p<0.01) decreased compared to pre-stroke (30.70±2.12 vs 48.89±2.35%, respectively, C). A significant (p<0.01) decrease in LF stride length was observed post-stroke compared to pre-stroke (59.04±3.85 vs 76.72±4.60cm, respectively, D). LF cycle time significantly (p<0.01) increased relative to pre-stroke (1.02±.09 vs 0.48±0.013sec, respectively, E). The mean pressure exhibited by the LF significantly (p<0.01) decreased at post-stroke compared to pre-stroke (2.62±.03 vs 2.82±.03 arbitrary units (A.U.), respectively, F). * indicates significant difference between pre-stroke and post-stroke time points.

## Discussion

In this study, we observed and characterized acute stroke injury severity, prognostic biomarkers, and potential therapeutic targets utilizing clinically relevant MRI, immune, behavior, and motor function tests in the translational ischemic stroke pig model. Lesion volumes were consistent among pigs and closely replicated human lesion volumes with similar impairments in functional performance (70–73). Ischemic injury produced cerebral swelling and consequent MLS as well as notable ICH, all of which are strongly associated with stroke patient morbidity (39, 74, 75). In addition, stroke led to reduced WM integrity of the IC correlating with a contralateral deteriorations in motor function commonly seen in patients post-stroke (30, 76, 77). Also similar to human stroke patients, MCAO led to an acute immune response marked by an increase in circulating neutrophils and a corresponding decrease in circulating lymphocytes which is a key biomarker for identifying ischemic stroke patients at risk for the development of intracranial hemorrhage thus influencing the use of tPA (78–80). Functional assessments showed impaired behavior and motor function disruptions that affected both spatiotemporal parameters and weight distribution, all of which parallel clinical functional outcomes in stroke patients (81–83). By further understanding these physiological hallmarks and exploiting the similarities between pigs and humans, the ischemic stroke pig model can be utilized to decrease the translational gap between rodent models and human stroke patients.

Early detection of ischemic infarction via DWI analysis has proven to be a critical component for both prognosis and therapeutic potentials within the narrow treatment window of acute ischemic stroke (84–86). This study showed mean lesion volumes of 9.91±3.14 cm^3^ at 24 hours post-stroke. Given that pig brains are approximately 7.5 times smaller than human brains, lesion volumes were found to closely replicate patient DWI lesion volumes. Acute DWI lesion thresholds of 72 cm^3^ are common in patients with major cerebral artery occlusions (26, 87–90). Often pre-clinical stroke models have relied on T1 or T2 MRI sequences which are typically delayed in early recognition of cerebral ischemia and do not account for diffusion abnormalities that may evolve into infarction (91–93). DWI lesion measurements overcome this limitation. Common pathological features of human ischemic infarction were also observed in our model including significant restricted diffusion in focal regions spanning the parietal, limbic, and temporal lobes as indicated by ADC maps (85, 94–96). Specifically, our pig model replicates characteristics of human MCA stroke by primarily demonstrating cytotoxic edema which will later evolve into vasogenic edema. In some pre-clinical stroke models, including rodent photochemical and photothromobotic models, cytotoxic edema and vasogenic edema develop simultaneously resulting in ischemic lesions lacking a penumbra (97, 98). This is a major model limitation as the penumbra is considered potentially salvagable tissue in human patients and is a coveted therapeutic intervention target. MRI-based discrimination of core from penumbra and non-ischemic tissue provides critical information for the testing of neuroprotective and restorative treatments as well as the initiation of surgical interventions within acute and sub-acute treatment windows (99, 100). For example, ischemic core volumes distinguishable from penumbra enable clinicians to consider the risk of cerebral hemorrhage from acute revascularization therapy (101). For these reasons, evaluating the efficacy and safety of potential treatments in an animal model with similar pathophysiology of acute ischemia in terms of cytotoxic and vasogenic edema as humans is of significant value.

Cerebral edema and consequent hemispheric swelling are serious stroke complications that result in rapid neurological deterioration and a disproportionately high 30-day patient mortality rate of 60-80% (102–104). Crudely managed via osmotic diuretics and/or decompressive craniectomies, patients are in desperate need for more effective and less invasive pharmacotherapies (105–107). These needs have been met with poor therapeutic translation due to discrepencies in lissencephalic small animal stroke models including limited cerebral edema and swelling as well as variable MLS and mortality rates (108–110). Specifically in endothelin-1 (ET-1) rodent stroke models, animals exhibit a dose-dependent ischemic lesion with marginal ischemic edema making this model less suited for studying acute stroke pathophysiology (111–114). In contrast, our pig stroke model exhibited increased ipsilateral hemipshere swelling due to the development of cytotoxic edema and consquent MLS within 24 hours post-stroke. These observations are in keeping with other large animal models of stroke, in which permanent ovine MCAO demonstrated cerebral edema and MLS (115). These physiological responses post-ischemic stroke are frequently associated with different levels of consciousness and serve as a predictive indicator of patient prognosis (116, 117). Furthermore, clinical studies indicate quantification of MLS can predict cerebral herniations and subsequent death prior to clinical signs and are a clinically relevant feature of this pig stroke model (118).

Although MRI techniques have become increasingly valuable in characterizing and refining the field’s understanding of ICH, the time course and underlying mechanisms remain poorly understood due to variability in the onset, size, and location of ICH in current stroke animal models (119). Often resulting from hemorrhagic transformation (HT) in ischemic stroke patients, spontaneous ICH incidence ranges from 38-71% in autopsy studies and from 13-43% in CT studies (120, 121). Furthermore, when ICH occupies >30% of the infarct zone, it has been correlated with early neurological deterioration and a significant increase in mortality rates 90 days post-ischemic stroke (122, 123). T2* sequences showed consistent mean hemorrhage volume between stroke pigs, indicating MCAO caused loss of macro- and microvessel integrity. The classical clinical presentations of ICH were replicated in our model through the progression of neurological deficits within hours post-stroke including decreased consciousness, head-pressing, vomiting, facial paralysis, and limb weakness (120, 124, 125). Interestingly, these neurological deficits correlated with the location of ICH. For example, ICH in the cerebellum was associated with ataxia whereas ICH in basal ganglia structures were associated with limb weakness. In previous studies, early neurologic deterioration was attributed primarily to cerebral edema and lesion volume; however, recent clinical pathological, MR, and CT studies suggest hemorrhage into ischemic tissues is a major contributor to poor clinical outcome, making ICH a novel target of pre-clinical studies (126–129). By replicating both tissue-level and neurological presentations unique to ICH, our model presents an exciting new platform for testing hemostatic therapies and surgical interventions.

For the first time, it was observed MCAO led to reduced WM integrity in the IC 24 hours post-stroke in the pig model. This subcortical structure is highly involved in communication between the cerebral cortex and brainstem resulting in profound muscle weakness and inhibited perception of sensory information of the patient’s face, arm, trunk, and leg post-stroke (130). Studies using Functional Ambulatory Categories found patients with IC lesions experience persistent (>6 months) functional motor deficits; requiring aids for balance and support during ambulation (131). As the right IC transmits nerve signals for movement of the left side of the body, our pig MCAO model closely replicated post-stroke deficits as seen through a decrease in spatiotemporal gait parameters of the hemiplegic limb including LF stride length and LF swing percent of cycle. Similarly, stroke patients exhibit decreases in the duration of stride length and the swing phase in the hemiplegic limb (132–135). Mean pressure of the LF limb was also decreased in stroke pigs likely as a result of overall greater weakness of the hemiplegic limb (136). Stroke pigs compensated for limb weakness and balance impairments by taking shorter, slower steps, thus reducing their velocity and cadence to better stabilize their gait. In a comparable human study utilizing the analogous GAITRite system, WM lesions corresponded with a poorer gait score as measured by step length and abnormal cadence in patients (77). These manifestations support our previous studies evaluating functional deficits post-stroke, thus providing further evidence quantitative gait analysis is a critical tool for the evaluation of stroke severity and therapeutic impact on recovery (25, 137).

Immune and inflammatory responses have been shown to play a key role in the sequela of ischemic stroke (138). Within the first few hours after stroke, neutrophils are recruited to the site of injury and release cytokines, chemokines, free oxygen radicals, and other inflammatory mediators (139). In this study, we observed a significant increase in neutrophils at 4 and 12 hours post-stroke. Neutrophil release of inflammatory mediators has been directly associated with cell damage or death as well as damage to the vasculature and extracellular matrices (139). Neutrophils have been implicated to play a significant role in blood brain barrier disruption and HT following ischemic stroke, which may explain one potential mechanism for HT observed 24 hours post-stroke in this study (79). Conversely, acute ischemic stroke has been shown to induce a rapid and long-lasting suppression of circulating immune cells such as lymphocytes that can lead to increased susceptibility of systemic infections after stroke (140). In this study, we observed a significant decrease in lymphocytes at 12 and 24 hours post-stroke, consistent with reports that stroke in humans induces immediate loss of lymphocytes that is most pronounced at 12 hours post-stroke (141). Though the exact mechanisms by which lymphocytes mediate immunosuppression post-stroke remain unclear, clinical evidence supports that lower levels of lymphocytes are a sign of poor long-term functional outcome (142–144). The neutrophil-to-lymphocyte ratio (NLR) was determined to be a useful marker to predict neurological deterioration and short-term mortality in patients with acute ischemic stroke (145, 146). Elevated NLRs have been reported to be associated with chronic inflammation, poor functional prognosis, and larger lesion volumes in ischemic stroke patients (78, 139, 146–148). These results suggest that neutrophil recruitment in our pig model may play a significant role in inflammatory-mediated secondary injury processes that contribute to the development of functional impairments. Furthermore, similar to human stroke patients, neutrophil and lymphocyte levels in our pig model may also serve as ideal markers for stroke severity and outcome prediction.

Open field testing is regularly used to evaluate behavior in rodents after ischemic stroke (149), specifically as an indicator of changes in exploratory behaviors (150, 151). In this study, a significant decrease was noted in perimeter sniffing frequency post-stroke in open field testing.

Pigs are inherently exploratory animals and perimeter sniffing is a typical exploratory behavior (152). This change in behavior may be attributed to post-stroke depression (PSD) as this behavioral disturbance has been reported to commonly develop in humans in the acute post-stroke period (153, 154). In accordance with the behavioral changes noted in the present study, PSD in humans is characterized by general apathy and lack of interest (155, 156). Evaluation and understanding of behavioral changes in a translational, large animal stroke model is crucial for future studies to assess functional outcomes of potential therapies.

In this study, we have demonstrated our pig model of ischemic stroke positively replicates cellular, tissue, and functional outcomes at acute time points similar to human stroke patients. MCAO in our pig ischemic stroke model exhibited a multifactorial effect leading to cytotoxic edema, lesioning, hemispheric swelling, and ICH, while also impairing diffusivity and WM integrity. These structural changes correlated with behavioral and motor function deficits in a similar manner to acute human stroke patients. As an effective model of acute ischemic stroke pathophysiology, the pig system is potentially an excellent tool for identifying potential treatment targets and testing novel therapeutics and diagnostics.

## Acknowledgements

The authors would like to thank Brandy Winkler and our team of undergraduate researchers who were involved in various aspects of surgeries, post-operative care, pig gait/behavioral testing, and data analysis. We would also like to thank the University of Georgia Animal Resources team for veterinary care and guidance as well as Rick Utley and Kelly Parham for their pig expertise and management skills.

